# Apelin-13 in the paraventricular nucleus protects myocardium from ischemia via V1a receptor of the paraventricular nucleus and GABAA receptor γ2 of the nucleus tractus solitarii in rats

**DOI:** 10.1101/2024.10.23.619957

**Authors:** Wen Yan, Dan Wang, Xinmin Zhang, Chengluan Xuan

**Affiliations:** Department of Anesthesia, The First Hospital of Jilin University, Jilin, China; Department of Anesthesia, The Second Hospital of Jilin University, Jilin, China

## Abstract

**Background:** Apelin system plays a significant role in central blood pressure regulation, but its role in neural control of myocardial injury protection is poorly understood. Thus, this study was undertaken to evaluate the effects of apelin-13 in the paraventricular nucleus (PVN) on myocardial infarction (MI) and ischemia/reperfusion (I/R).

**Methods:** The cardiac functions were assessed after microinjecting or transferring the apelin-13 gene into the paraventricular nucleus (PVN) of rat models with myocardial infarction (MI) or ischemia/reperfusion (I/R).

**Results:** In MI rats, we showed that apelin-13 expression decreased in PVN, V1a receptor expression increased in PVN and nucleus tractus solitarii (NTS), and GABAA receptor (GAR) γ2 expression increased in NTS. In primary cultured hypothalamus and medulla oblongata neurons, V1a receptor expression was downregulated by APJ receptor antagonist or GAR agonist, indicating that there are interactions between these receptors in neurons. Apelin-13 overexpression in PVN significantly improved cardiac function of MI and I/R rats, including left-ventricular end-diastolic diameter, left-ventricular end-systolic diameter, left-ventricular ejection fraction, and left-ventricular fractions shortening, accompanied by decreased noradrenaline and increased vasopressin plasma levels. Myocardial ischemia–related apoptotic and inflammatory pathway markers (Bcl-1, Bax, TGF-β1, CCR5, Smad2) were downregulated and four neuropeptides of the parasympathetic endocrine system (somatostatin, cholecystokinin, glucagon-like peptide 1, vasoactive intestinal peptide) were increased in serum correspondingly with cardiac function improving. V1a receptor antagonist in PVN or NTS and GAR agonist in NTS decreased the effects of apelin-13 overexpression on cardiac function of MI and I/R rats.

**Conclusions:** Overexpression of apelin-13 in PVN leads to cardiac function improvement via V1a receptor in PVN and NTS, and GARγ2 in NTS. Parasympathetic endocrine system and myocardial ischemia–related apoptotic and inflammation signaling pathways are involved in apelin-13 in PVN–mediated cardiac function regulation, which provides evidence for neural regulation of cardiovascular diseases.

## Introduction

Brain–heart crosstalk is now referred to as neurocardiology, which addresses the effects of brain injury on the heart or the effects of cardiac injury on the brain.^1^ There is increasing experimental and clinical evidence not only suggesting a causal link from the heart to the brain, but also indicating the presence of a bidirectional relationship, highlighting a powerful influence of the brain on the heart.^2^

Activation of the hypothalamus–pituitary–adrenal (HPA) axis plays a pivotal role in communication between brain and heart leading to cardiac dysfunction.^3^ The brain can exhibit a variety of effects on cardiac function through signaling via the sympathetic nervous system, parasympathetic nervous system, renin–angiotensin– aldosterone system, and HPA axis.^4^ The paraventricular nucleus (PVN) of the hypothalamus acts as the primary integrator that modulates HPA axis activity, which can cause increased blood pressure, alter heart rate, and promote inflammation in the cardiovascular system.^5^ The PVN is also a master controller of the autonomic nervous system, which provides specialized innervation to all autonomic relay centers^6^ and together with the dorsomedial hypothalamic nucleus, it integrates neuroendocrine, homeostatic, and stress response.^7^ The medullary nucleus tractus solitarii (NTS) is reciprocally connected with the pontine parabrachial and Kölliker–Fuse nuclei to relay visceral afferent information to other central autonomic network (CAN) structures.^8^ The NTS relays afferent information to the PVN, which sends efferent projections to the rostral lateral medulla (RVLM) and spinal cord to regulate sympathetic outflow in turn. Reduction of the PVN’s activity promotes improved recovery of cardiac function after myocardial infarction.^9^

Apelin, containing 77 amino acids, has been isolated from bovine stomach extracts, and its effects are mediated via ligand for the orphan G-protein–coupled (APJ) receptor.^10, 11^ Apelin-13 is the main isoform that activates the APJ receptor. As a novel activity regulator, the apelin/APJ system is involved in neurohumoral regulation in the hypothalamus–pituitary axis^12^ and is widely expressed in microglia and neurons of the CNS with emerging evidence.^13^ Microinjection of apelin-13 in the PVN induces long-term high blood pressure and sympathetic activity mediated by superoxide anions.^14^ Besides that, the apelin/APJ system in the RVLM also plays important roles in central blood pressure regulation and pathogenesis of hypertension.^15^ Based on previous data, apelin-13 may be considered to affect the cardiovascular system via the vasopressinergic system in the PVN and supraoptic nucleus (SON) of the hypothalamus.^16, 17^ In the hypothalamus and medulla oblongata, vasopressin effects are primarily mediated through V1a receptors to perform the central regulation of the cardiovascular system.^18^

γ-Aminobutyric acid (GABA) is a well-known neuronal transmitter that exerts inhibitory actions in the brain mediated by GABA_A_ receptor (GAR) and GABA_B_ receptor (GBR), which are defined based on pharmacological and physiological research.^19^ According to our previous study, GABAergic system in the NTS contributes to central resetting of blood pressure and development of hypertension.^20^ It has been indicated that baroreceptor activation could inhibit vasopressin secretion, which might be regulated by the activation of GABAergic projections to vasopressin neurons, and local administration of GAR antagonist could inhibit vasopressin neurons during the onset of hypertension.^21^ Thus, the apelin/APJ system, vasopressinergic system, and GABAergic system are vital pathways of the CNS involved in the neural control of the cardiovascular system in the hypothalamus, but they remain poorly understood in relation to myocardial injury. The reduction in APJ mRNA expression has been observed in the hypothalamus in post-infarct heart failure, especially in the presynaptic neurons of the PVN, which could attenuate the pressor response to intracerebroventricularly infused apelin-13.^22^

The parasympathetic endocrine system (PES) comprises circulating peptides released from secretory cells that are significantly modulated by vagal projections from the dorsal motor nucleus of the vagus nerve (DMV).^23^ Four peptides, including somatostatin (SST), cholecystokinin (CCK), glucagon-like peptide 1 (GLP-1), and vasoactive intestinal peptide (VIP), presented in the circulatory system perform sympathetic antagonism to balance sympathetic activation–induced pupillary dilation and increased rate and contractility of the heart.^24^

In this study, we aimed to assess the effects of apelin-13 in the PVN on cardiac function of rats with myocardial infarction (MI) and ischemia/reperfusion (I/R). The involved mechanisms were also explored, including myocardial ischemia–related apoptotic and inflammatory pathway markers in heart and neuropeptides in serum.

## Methods and materials

All animal procedures were conducted in accordance with the institutional guidelines of The First Hospital of Jilin University and were approved by the Institutional Animal Care and Use Committee (2020-0128). All studies were approved locally and conformed to the guidelines from Directive 2010/63/EU of the European Parliament on the protection of animals used for scientific purposes. The Sprague–Dawley rats were purchased from Charles River. The rats were maintained under a 12/12 h light/dark cycle with food and water provided by The First Hospital of Jilin University. The rat studies were performed under isoflurane anesthesia (2.0%–2.5% isoflurane, 100% oxygen at 2 L/minute) in spontaneous breathing. To perform histology studies and blood tests, sodium pentobarbital (40–60 mg/kg) was intraperitoneally injected to anesthetize the rats. The rats were then killed by exsanguination while still under anesthesia. The neonatal rats for primary neuron cultures were killed by overdose of intraperitoneally injected sodium pentobarbital. Other details and expanded protocols are described in the Supplementary Material. The animal experiments were carried out as shown in Supplementary Fig. 1.

## Results

### Experiments in rats with coronary artery ligation

1. Expression of apelin-13, V1a receptor, and GARγ2 receptor in MI rats: On day 28 after MI, apelin-13 expression was significantly lower at both protein and mRNA levels in the PVN, but not in the NTS and RVLM, compared with the sham rats (Fig. 1A–C; n=5). V1a receptor expression was increased in the PVN and NTS, but not in the RVLM at both protein and mRNA levels (Fig. 1A, D, E; n=5). GARγ2 was increased in the NTS, but not in the PVN and RVLM in the MI rats compared with the sham control. In contrast, GBR1 expression was not significantly affected (Fig. 1A, F, G).
2. Effects of apelin-13 microinjection into the PVN on cardiac function in MI rats: Echocardiography on day 28 after MI showed improved cardiac function in the rats receiving continuous apelin-13 microinjection into the PVN. Specifically, these rats showed higher left-ventricular ejection fraction (LVEF) (62.7%±3.0% vs. 51.0±3.2%, *p*<0.01) (Fig. 1J) and higher left-ventricular fractions shortening (LVFS) (38.3%±1.5% vs. 26.4±2.1%, *p*<0.001) (Fig. 1K), and lower left-ventricular end-diastolic diameter (LVEDD) (6.34±0.45 vs. 8.06±0.16 mm, *p*<0.01) (Fig. 1L) and lower left-ventricular end-systolic diameter (LVESD) (5.97±0.20 mm vs. 7.74±0.24 mm, *p*<0.001) compared with the MI group (Fig. 1M). Continuous apelin-13 microinjection into the PVN in the MI rats increased V1a receptor expression in the PVN and NTS, but decreased GARγ2 expression in the NTS (Fig. 1N–P). Compared with the MI rats, the MI rats receiving apelin-13 microinjection into the PVN had lower serum levels of noradrenaline (578.3±51.5 vs. 825.0±89.4 pg/mL), but higher levels of vasopressin (18.9±0.8 vs. 11.5±0.8 pg/mL) (Fig.1U–V).
3. Effects of apelin-13 microinjection into the PVN on neuropeptides: Compared with MI rats, the MI rats receiving apelin-13 microinjection into the PVN had higher serum levels of SST (131.7±6.6 vs. 79.8±4.5 pg/mL, *p*<0.001), CCK (52.7±5.6 vs. 25.6±1.1 pg/100 mL, *p*<0.001), VIP (163.8±8.3 vs. 105.6±6.3 pg/mL, *p*<0.001), and GLP-1 (12.4±0.7 vs. 10.0±1.1 pg/mL, *p*<0.05) (Fig. 1Q–T).
4. Expression of myocardial ischemia–related apoptotic proteins after apelin-13 overexpression (AAV2–apelin-13 gene transfer) in the PVN of the MI rats: Overexpression of apelin-13 in the PVN (Fig. 2Aa–Af) of the MI rats resulted in a time-dependent increase in Bcl-2 and a decrease in Bax expression in the heart (Fig. 2B, C–D). TGF-β1, CCR5, and Smad2 decreased over time (Fig. 2B, E–G). V1a receptor expression in the PVN and NTS increased, and GARγ2 expression in the NTS decreased gradually over time (Fig. 2B, H–J).
5. Effects of APJ receptor and GARγ2 receptor on V1a receptor in primary cultured neurons of the hypothalamus and medulla oblongata: Immunofluorescence staining in a primary culture of hypothalamus and medulla oblongata neurons (Fig. 3A–D) showed decreased V1a receptor expression at both protein (Fig. 3E, F) and mRNA levels (Fig. 3G) upon treatment with either APJ receptor antagonist F13A (Fig. 3A) or the GABA-A agonist muscimol (Fig. 3B) in a time-dependent manner.
6. Effect of apelin-13 overexpression (AAV2–apelin-13 gene transfer) in the PVN on V1a receptor–mediated cardiac function improvement of the MI rats: Overexpression of apelin-13 in the PVN of the MI rats significantly improved cardiac function, including LVEF (79.1%±1.8% vs. 66.2%±2.2%, *p*<0.05) (Fig. 4A), LVFS (45.0%±1.9% vs. 32.8%±1.3%, p<0.05) (Fig. 4B), LVEDD (7.14±0.45 vs. 8.34±0.36 mm, *p*<0.05) (Fig. 4C), and LVESD (5.01±0.17 vs. 6.12±0.23 mm, *p*<0.05) (Fig. 4D). These effects were attenuated by continuous microinjection of the V1a receptor antagonist SR49059 into the PVN and NTS and continuous microinjection of the GABA-A antagonist muscimol into the NTS (Fig. 4A–D). Compared with continuous microinjection of SR49059 into the PVN of the MI rats with apelin-13 overexpression in the PVN, continuous microinjection of SR49059 or muscimol into the NTS had less attenuation effects (Fig. 4A–D). Protective effects of apelin-13 overexpression in HE (Fig. 4E), Masson (Fig. 4F), TUNEL (Fig. 4G), and TTC (Fig. 4H) were evident after microinjection of SR49059 into the PVN or NTS, and after muscimol injection into the NTS. Bar graphs of Masson staining, TUNEL, and TTC staining indicated that apelin-13 overexpression in the PVN significantly decreased fibrosis (29.0%±3.4% vs. 56.0%±3.4%, *p*<0.05) (Fig. 4I), mean fluorescence intensity (27.27±2.81 vs. 55.34±3.32 AU, *p*<0.05) (Fig. 4J), and infarction area (24.4%±4.1% vs. 46.8%±4.9%, *p*<0.05) (Fig. 4K). Compared with SR49059 microinjection into the PVN, SR49059 or muscimol microinjection into the NTS led to less fibrosis, lower mean fluorescence intensity, and reduced infarction area (Fig. 4I–K). Compared with the sham group, the MI rats with apelin-13 overexpression in the PVN had similar MAP (94.9±2.7 vs. 97.1±4.1 mm Hg, n=7, *p*>0.05) and HR (383.4±9.1 vs. 395.4±10.7 bpm, n=7, *p*>0.05) (Fig. 4L–M) at 28 days after MI model surgery. Increased V1a receptor expression in the PVN by AAV2-apelin-13 gene overexpression in the PVN was attenuated by continuous microinjection of the V1a receptor antagonist SR49059 into the NTS, but not into the PVN or the GABA-A antagonist muscimol microinjection into the NTS (Fig. 5A, B). In the NTS, increased V1a receptor expression by AAV2-apelin-13 gene overexpression in the PVN was not attenuated by continuous microinjection of the V1a receptor antagonist SR49059 into the PVN or NTS, or by the GABA-A antagonist muscimol into the NTS (Fig. 5A, C). Decreased GARγ2 expression in the NTS by AAV2-mediated apelin-13 overexpression in the PVN was significantly attenuated by continuous microinjection of the V1a receptor antagonist SR49059 in the NTS, but not by microinjection of SR49059 in the PVN or muscimol in the NTS (Fig. 5A, D). Apelin-13 gene overexpression–induced increase in Bcl-2 (Fig. 5E, F) and decrease in Bax (Fig. 5E, G), TGF-β1 (Fig. 5E, H), CCR5 (Fig. 5E, I), and Smad2 (Fig. 5E, J) were attenuated by microinjection of SR49059 into the PVN, but not into the NTS or muscimol injection into the NTS. However, in serum, increased levels of SST (Fig. 5K), CCK (Fig. 5L), VIP (Fig. 5M), GLP-1 (Fig. 5N), and vasopressin (Fig. 5P) induced by apelin-13 overexpression in the PVN of the MI rats were attenuated by SR49059 in the PVN or NTS and muscimol in the NTS. Contrarily, decreased level of noradrenaline was enhanced by SR49059 in the PVN or NTS and muscimol in the NTS (Fig. 5O).

**Fig. 1.**
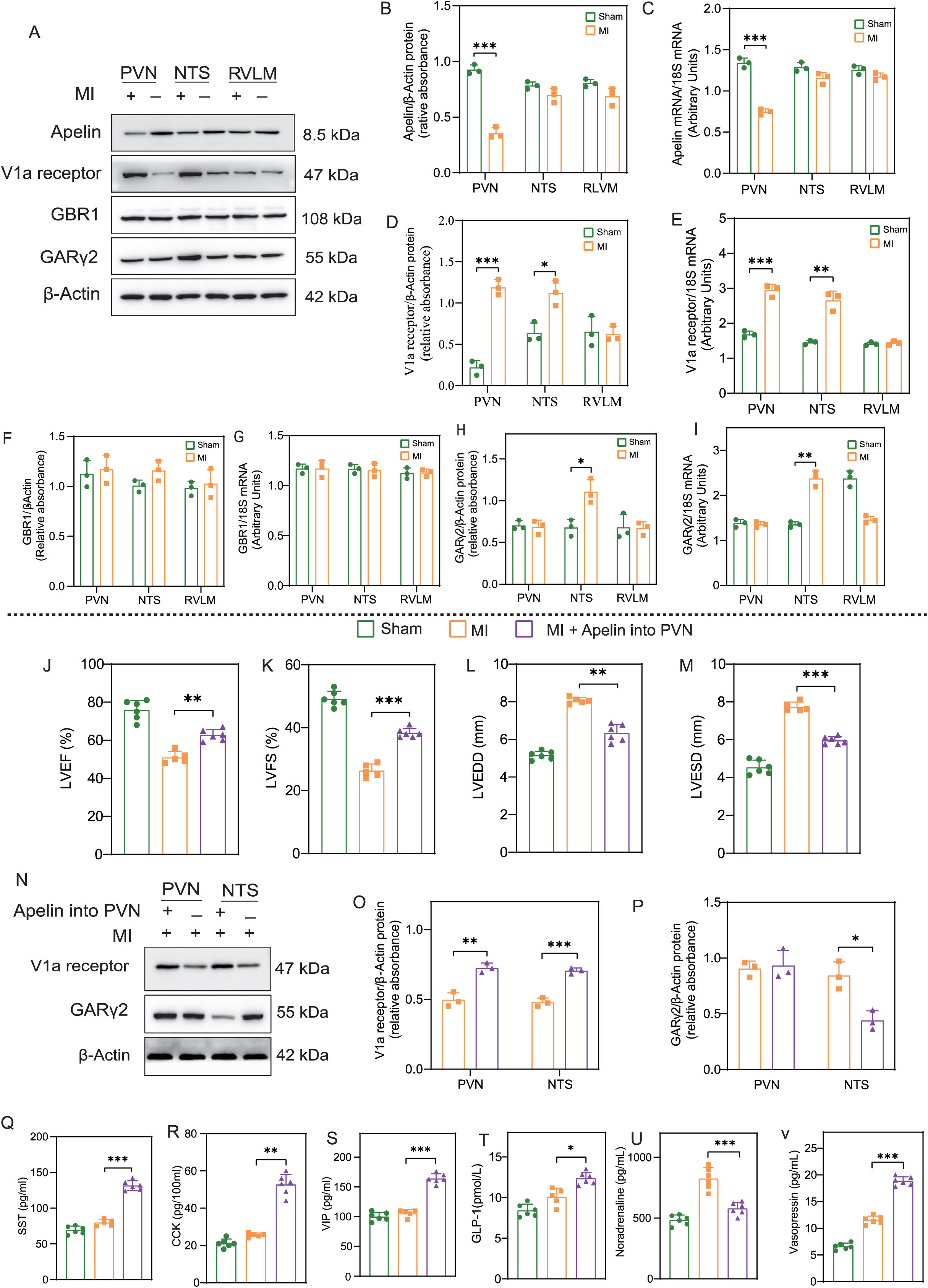
Expression of apelin-13, V1a receptor, and GARγ2 receptor, and effects of apelin-13 microinjection into the PVN on cardiac function in MI rats. **A** Representative western blot (WB) for apelin-13, V1a receptor, GBR1 receptor, and GARγ2 receptor in the PVN, NTS, and RVLM from MI rats and sham-surgery rats. **B** Quantification of apelin-13 protein levels. **C** mRNA expression levels of apelin-13. **D** Quantification of V1a receptor protein levels. **E** mRNA expression levels of V1a receptor. **F** Quantification of GBR1 receptor protein levels. **G** mRNA expression levels of GBR1. **H** Quantification of GARγ2 receptor protein levels. **I** mRNA expression levels of GARγ2. **J–M** Echocardiographic results of LVEF, LVFS, LVEDD, and LVESD (n=6 in the sham group; n=5 in the MI group, 1 rat lost; n=6 in the MI + Apelin group). **N** Representative WB for V1a receptor and GARγ2 in the PVN and NTS from MI rats and MI rats with apelin-13 continuously microinjected into the PVN for 28 days. **O** Quantification of V1a receptor protein levels. **P** Quantification of GARγ2 receptor protein levels. **Q–V** The plasma levels of SST, CCK, VIP, GLP-1, noradrenaline, and vasopressin (n=6 in the sham group; n=5 in the MI group, 1 rat lost; n=6 in the MI + Apelin group). Normality was tested using the Shapiro–Wilk test. The data (n=3 experiments) in **B**–**I** are shown as mean ± SD and were analyzed using unpaired two-tailed *t* test. The data in **J**–**M** are shown as mean ± SD, and were analyzed using one-way ANOVA, corrected using Tukey’s multiple-comparisons test. The data (n=3 experiments) in **O**–**P** are shown as mean ± SD and were analyzed using unpaired two-tailed *t* test. The data in **Q**–**V** are shown as mean ± SD, and were analyzed using one-way ANOVA, corrected using Tukey’s multiple-comparisons test. *, *p*<0.05; **, *p*<0.01; ***, *p*<0.001.

**Fig. 2.**
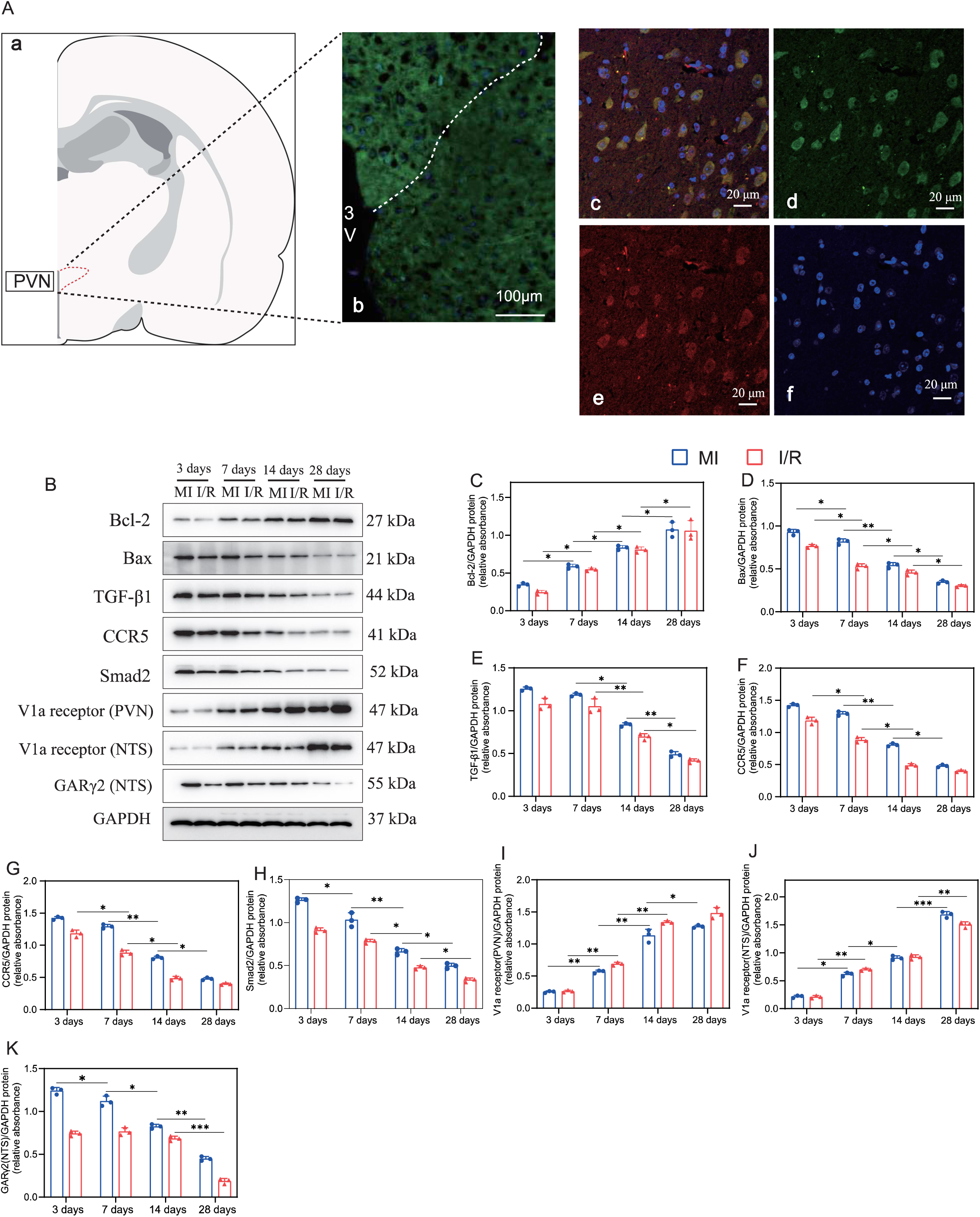
Expression of myocardial ischemia–related apoptotic proteins after apelin-13 overexpression in the PVN of MI or I/R rats. **A** Immunofluorescence images showing overexpression of apelin-13 protein in the PVN of MI rats after 28 days of apelin gene transfer (n=6 in each group). **a**) Location of the stained PVN brain sections shown in b–f, based on the rat brain atlas of Swanson.^38^ **b**) Fluorescence micrograph (×10 magnification) demonstrating localization of apelin-13 within the PVN. 3v is the third ventricle. **c**) Overlap of **d**, **e**, and **f**, showing that the green fluorescence is neuronally located. **d**) Fluorescence micrograph (×40 magnification) indicating localization of apelin-13 in PVN cells, immunostained with anti-apelin antibody (green). **e**) Same filed of cells as in panel **d**, immunostained with anti-NeuN antibodies (red). **f**) DAPI of the same filed of cells in panel **d**. **B** Molecular pathways regulating apoptotic and inflammation pathways were determined through WB for TGF-β, CCR-5, Smad2, Bax, and Bcl-2 in time course of I/R or MI rats, and expression of V1a receptor and GARγ2 induced by apelin-13 overexpression in the PVN. **C–J** Quantification of fold change vs. GAPDH. The data (n=3 experiments) in **C**–**J** are shown as mean ± SD, and were analyzed using unpaired two-tailed *t* test. *, *p*<0.05; **, *p*<0.01; ***, *p*<0.001.

**Fig. 3.**
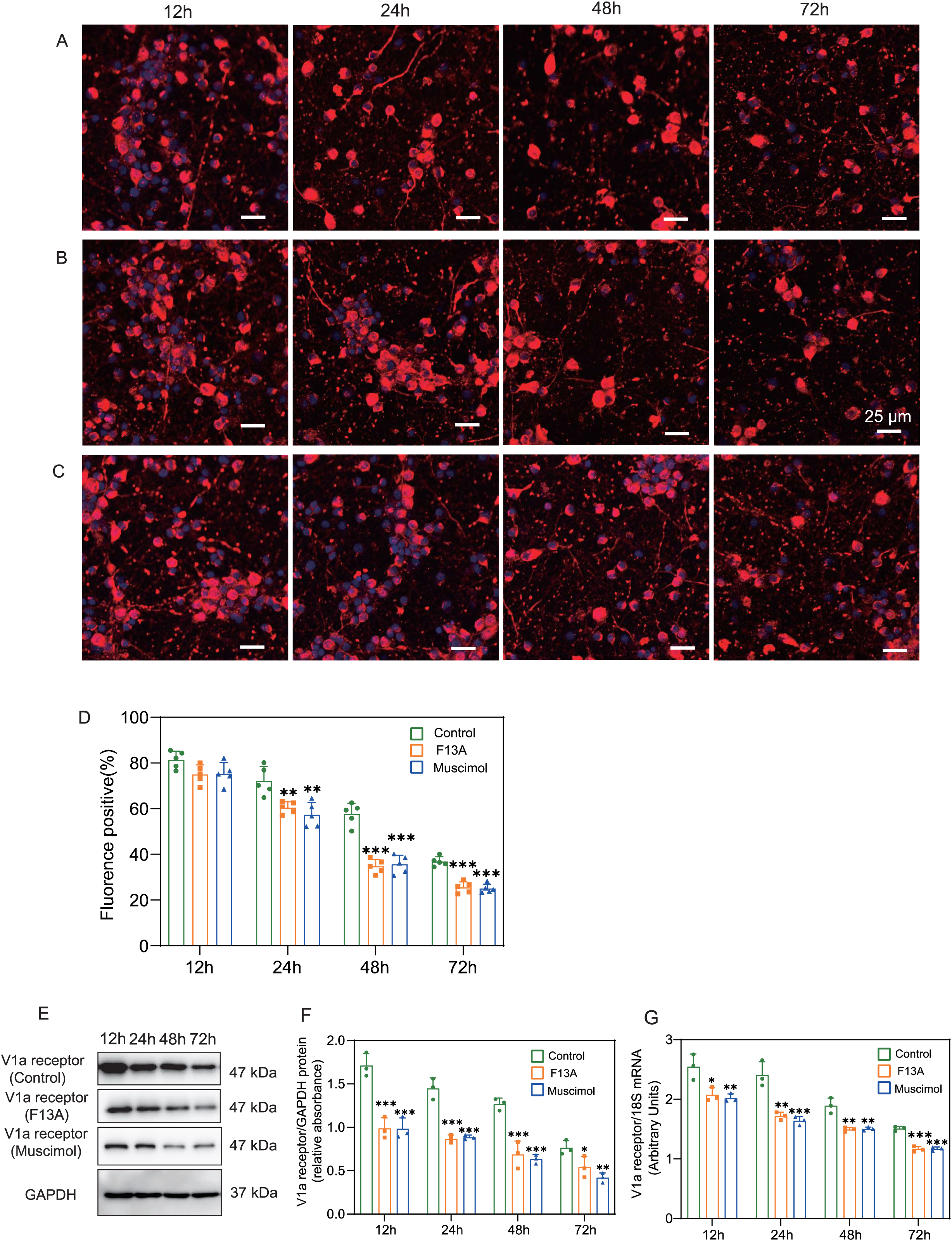
Effects of APJ receptor and GARγ2 receptor on V1a receptor in primary cultured neurons of the hypothalamus and medulla oblongata. **A–C** Immunofluorescence for F13A-, muscimol-, and placebo-treated primary neurons of the hypothalamus and medulla oblongata. Neurons treated for 12, 24, 48, and 72 hours. **D** Representative positive fluorescence. **E** Representative WB for V1a receptor in primary cultured neurons treated with F13A and muscimol, and **F** its quantification. **G** mRNA expression levels of V1a receptor in neurons treated with F13A and muscimol. The data (n=5 experiments) in **D** are shown as mean ± SD, and were analyzed using one-way ANOVA, corrected using Dunett’s multiple-comparisons test. The data (n=3) in **F**–**G** are shown as mean ± SD, and were analyzed using one-way ANOVA, corrected using Dunett’s multiple-comparisons test. *, *p*<0.05; **, *p*<0.01; ***, *p*<0.001, compared with the control group.

**Fig. 4.**
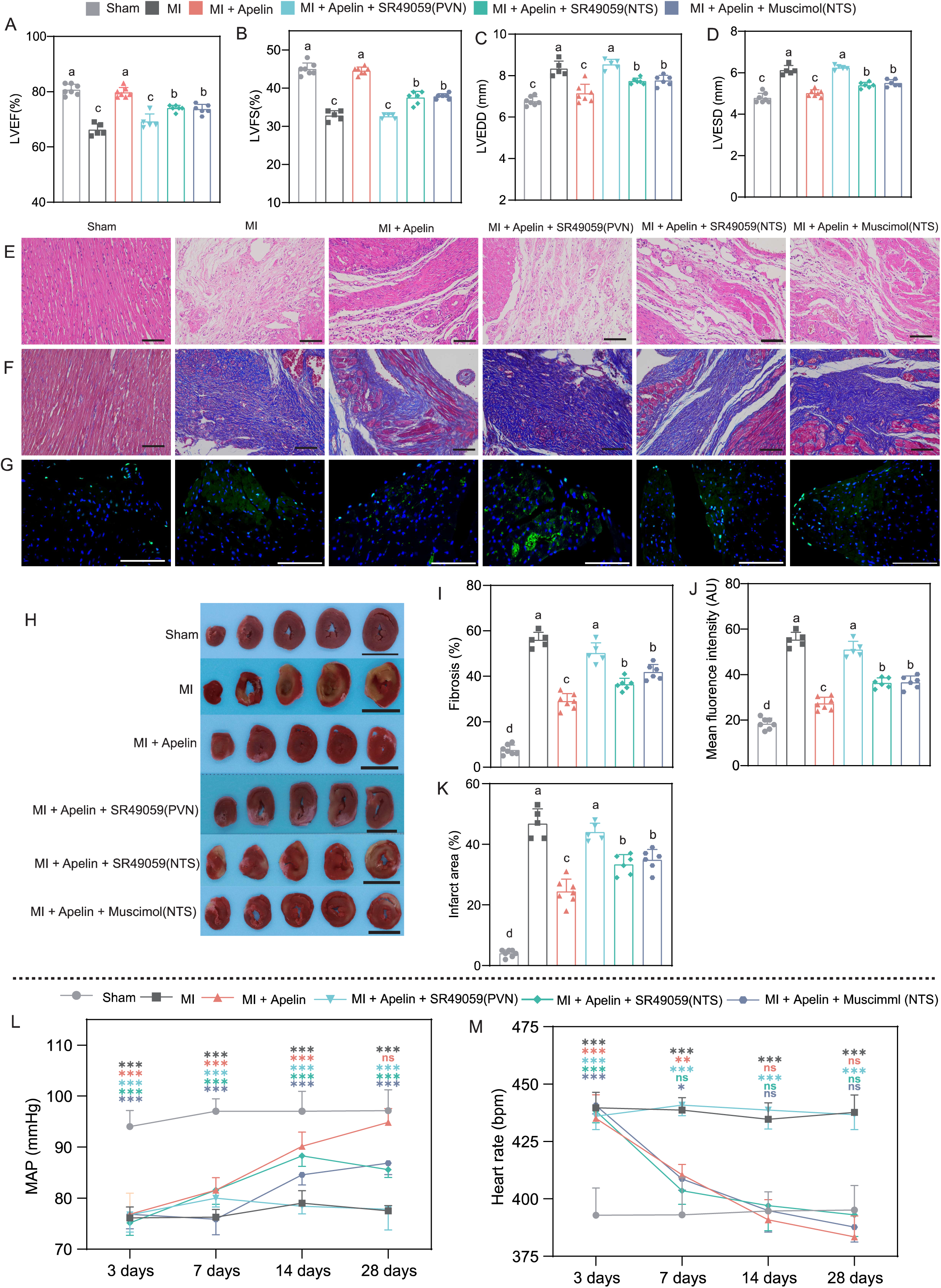
Effect of apelin-13 overexpression (AAV2-apelin-13 gene transfer) in the PVN on V1a receptor–mediated cardiac function and cardiac morphological improvement of MI rats. **A–D** Echocardiographic results of LVEF, LVSF, LVEDD, and LVESD. **E** Representative HE staining. **F** Representative Masson staining and **I** its quantification. **G** Representative TUNEL staining and **J** its quantification. **H** Representative TTC staining and **K** its quantification. Black bar in E and f is 100 μm; white bar in G is 50 μm; black bar in H is 1 cm. At 28 days after MI model surgery, the sham group had 7 rats, the MI group had 5 rats (2 rats lost), the MI + Apelin group had 7 rats, the MI + Apelin + SR49059 (PVN) group had 5 rats (2 rats lost), the MI + Apelin + SR49059 (NTS) group had 6 rats (one rat lost), and the MI + Apelin + Muscimol (NTS) group had 6 rats (one rat lost). Normality was tested using the Shapiro–Wilk test. Data in **A**–**D** and **I**–**M** are shown as mean ± SD, and were analyzed using one-way ANOVA, corrected using Tukey’s multiple-comparisons test. In **A**–**D** and **I**–**K**, the differences are indicated by the abc annotation method. MAP (mean arterial pressure) (**L**) and heart rate (**M**) were recorded in a conscious state at 3 days, 7 days, 14 days, and 28 days after MI model surgery. *, *p*<0.05; **, *p*<0.01; ***, *p*<0.001; ns, not significant, compared with the sham group.

**Fig. 5.**
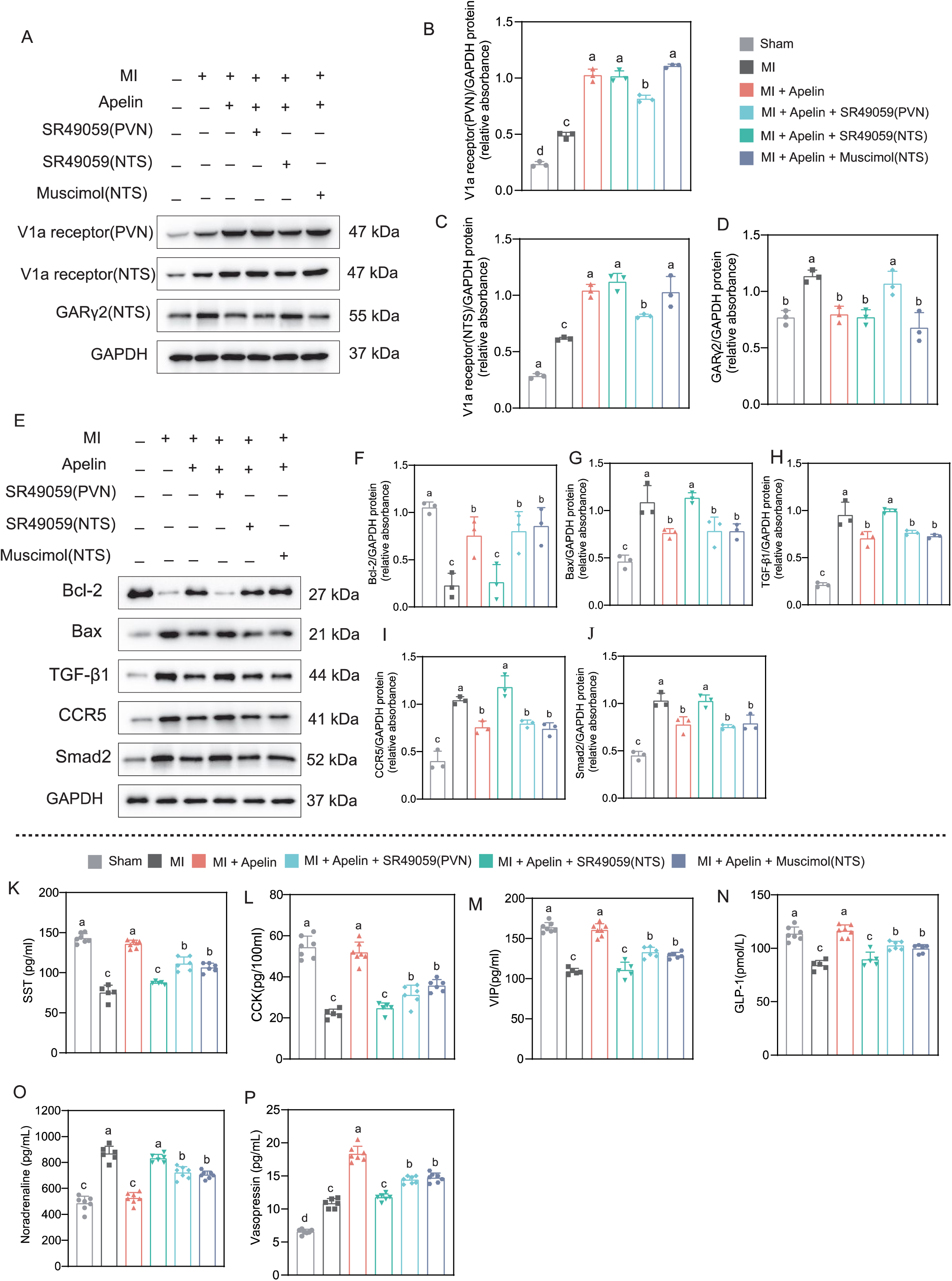
Mechanisms of the effect of apelin-13 overexpression (AAV2-apelin-13 gene transfer) in the PVN on V1a receptor–mediated cardiac function improvement of MI rats. **A** Representative WB for V1a receptor in the PVN or NTS and GARγ2 expression in the NTS, and **B–D** its quantification. **E** Representative WB for Bcl-2, Bax, TGF-β1, CCR-5, and Smad2 protein level in the heart and **F–J** its quantification. **K–N** The plasma levels of SST, CCK, VIP, and GLP-1 (n=7 in sham group; n=5 in the MI group, 2 rats lost; n=7 in the MI + Apelin group; n=5 in the MI + Apelin + SR49059 (PVN) group, 2 rats lost; n=6 in the MI + Apelin + SR49059 (NTS) group, one rat lost; n=6 in the MI + Apelin + Muscimol (NTS) group, one rat lost). **O** and **P** Representative plasma levels of noradrenaline and vasopressin (n=7 in the sham group; n=6 in the MI group, one rat lost; n=7 in the MI + Apelin group; n=6 in the MI + Apelin + SR49059 (PVN) group, one rat lost; n=6 in the MI + Apelin + SR49059 (NTS) group, one rat lost; n=7 in the MI + Apelin + Muscimol (NTS) group). Normality was tested using the Shapiro–Wilk test. Data (n=3 experiments) in B–D and F–J, and data in K–P are shown as mean ± SD, and were analyzed using one-way ANOVA, corrected using Tukey’s multiple-comparisons test. In **B**–**D** and **F**–**P**, the differences are indicated by the abc annotation method.

## Experiments in rats with ischemia followed by reperfusion

1. Expression of myocardial ischemia–related apoptotic proteins after apelin-13 overexpression in the PVN of I/R rats: Overexpression of apelin-13 in the PVN using an AAV2 in the I/R rats resulted in a time-dependent increase in Bcl-2 and a decrease in Bax expression in the heart (Fig. 2B, C–D). TGF-β1, CCR5, and Smad2 also decreased over time (Fig. 2B, E–G). V1a receptor expression in the PVN and NTS increased, and GARγ2 expression in the NTS decreased gradually over time (Fig. 2B, H–J). These results were similar to those of apelin-13 overexpression in the PVN of the MI rats.
2. Effect of apelin-13 overexpression (AAV2-apelin-13 gene transfer) in the PVN on V1a receptor–mediated cardiac function improvement of the I/R rats: Overexpression of apelin-13 in the PVN of the I/R rats significantly improved cardiac function, including LVEF (80.4%±1.5% vs. 72.9%±2.3%, *p*<0.05) (Fig. 6A), LVFS (44.6%±1.8% vs. 37.7±1.4%, *p*<0.05) (Fig. 6B), LVEDD (7.10±0.24 vs. 8.22±0.19 mm, *p*<0.05) (Fig. 6C), and LVESD (5.09±0.20 vs. 6.26±0.16 mm, *p*<0.05) (Fig. 6D). Continuous microinjection of SR49059 into the PVN or NTS and microinjection of muscimol into the NTS attenuated the effects on LVEF and LVFS, but SR49059 or muscimol in the NTS did not attenuate the effects on LVEDD and LVESD (Fig. 6A–D). Protective effects of apelin-13 overexpression in HE (Fig. 6E), Masson (Fig. 6F), TUNEL (Fig. 6G), and TTC staining (Fig. 6H) were evident by microinjection of SR49059 into the PVN or NTS and muscimol injection into the NTS. Bar graphs of Masson, TUNEL, and TTC staining indicated that apelin-13 overexpression in the PVN of the I/R rats significantly decreased fibrosis (30.8%±1.7% vs. 50.9%±3.3%, *p*<0.05) (Fig. 6I), mean fluorescence intensity (24.77±1.77 vs. 78.45±5.49 AU, *p*<0.05) (Fig. 6J), and infarction area (8.6%±1.8% vs. 19.0%±1.8%, *p*<0.05) (Fig. 6K), which was attenuated by SR49059 microinjection into the PVN. Compared with SR49059 microinjection into the PVN, SR49059 or muscimol microinjection into the NTS led to less fibrosis, lower mean fluorescence intensity, and reduced infarction area (Fig. 6I–K). Compared with the sham group, the I/R rats with apelin-13 overexpression in the PVN had similar MAP (93.9±1.9 vs. 96.3±4.4 mm Hg, n=7, *p*>0.05) and HR (384.0±9.0 vs. 386.4±6.0 bpm, n=7, *p*>0.05) (Fig. 4L–M) at 28 days after I/R model surgery. Increased V1a receptor expression in the PVN induced by apelin-13 overexpression in the PVN was attenuated by continuous microinjection of SR49059 into the NTS, but not into the PVN or by injection of muscimol into the NTS (Fig. 7A, B). In the NTS, increased V1a receptor expression by apelin-13 overexpression in the PVN was not attenuated by continuous microinjection of SR49059 into the PVN or muscimol into the NTS, but attenuated by SR49059 into the NTS (Fig. 7A, C). Decreased GARγ2 expression in the NTS induced by AAV2-mediated apelin-13 overexpression in the PVN was significantly attenuated by continuous microinjection of SR49059 into the NTS, but not into the PVN, or injection of muscimol into the NTS (Fig. 7A, D). Apelin-13 overexpression– induced increase in Bcl-2 (Fig. 7E, F) and decreases in Bax (Fig. 7E, G), TGF-β1 (Fig. 7E, H), CCR5 (Fig. 7E, I), and Smad2 (Fig. 7E, J) were attenuated by microinjection of SR49059 into the PVN, but not into the NTS, or by muscimol injection into the NTS. However, in serum, increased levels of SST (Fig. 7K), CCK (Fig. 7L), VIP (Fig. 7M), GLP-1 (Fig. 7N), and vasopressin (Fig. 7P) induced by apelin-13 overexpression in the PVN of the MI rats were attenuated by SR49059 in the PVN or NTS and muscimol in the NTS. Conversely, the decreased level of noradrenaline was enhanced by SR49059 in the PVN and muscimol in the NTS (Fig. 7O). Thus, the effect of apelin-13 overexpression in the PVN on V1a receptor showed similar cardiac functions improvement for the MI and I/R rats.

**Fig. 6.**
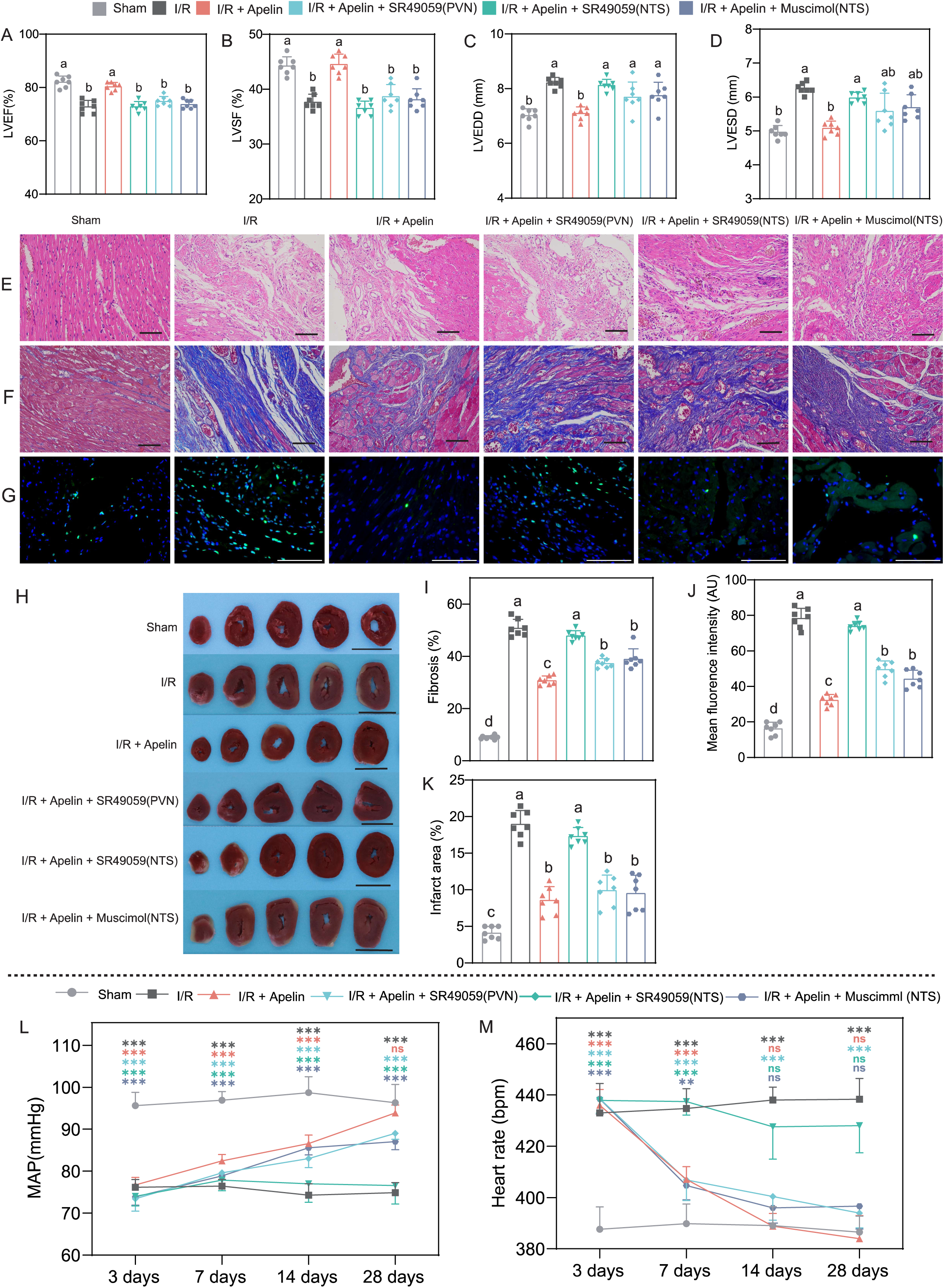
Effect of apelin-13 overexpression (AAV2-apelin-13 gene transfer) in the PVN on V1a receptor–mediated cardiac function and cardiac morphological improvement of I/R rats. **A–D** Echocardiographic results of LVEF, LVSF, LVEDD, and LVESD. **E** Representative HE staining. **F** Representative Masson staining and **I** its quantification. **G** Representative TUNEL staining and **J** its quantification. **H** Representative TTC staining and **K** its quantification. Black bar in **E** and **F** is 100 μm; white bar in **G** is 50 μm; black bar in **H** is 1 cm. At 28 days after I/R model surgery, each group had 7 rats. Normality was tested using the Shapiro–Wilk test. Data in A–D and I–K are shown as mean ± SD, and were analyzed using one-way ANOVA, corrected using Tukey’s multiple-comparisons test. In **A**–**D** and **I**–**K**, the differences are indicated by the abc annotation method. MAP (mean arterial pressure) (**L**) and heart rate (**M**) were recorded in a conscious state at 3 days, 7 days, 14 days, and 28 days after I/R model surgery. *, *p*<0.05; **, *p*<0.01; ***, *p*<0.001; ns, not significant, compared with the sham group.

**Fig. 7.**
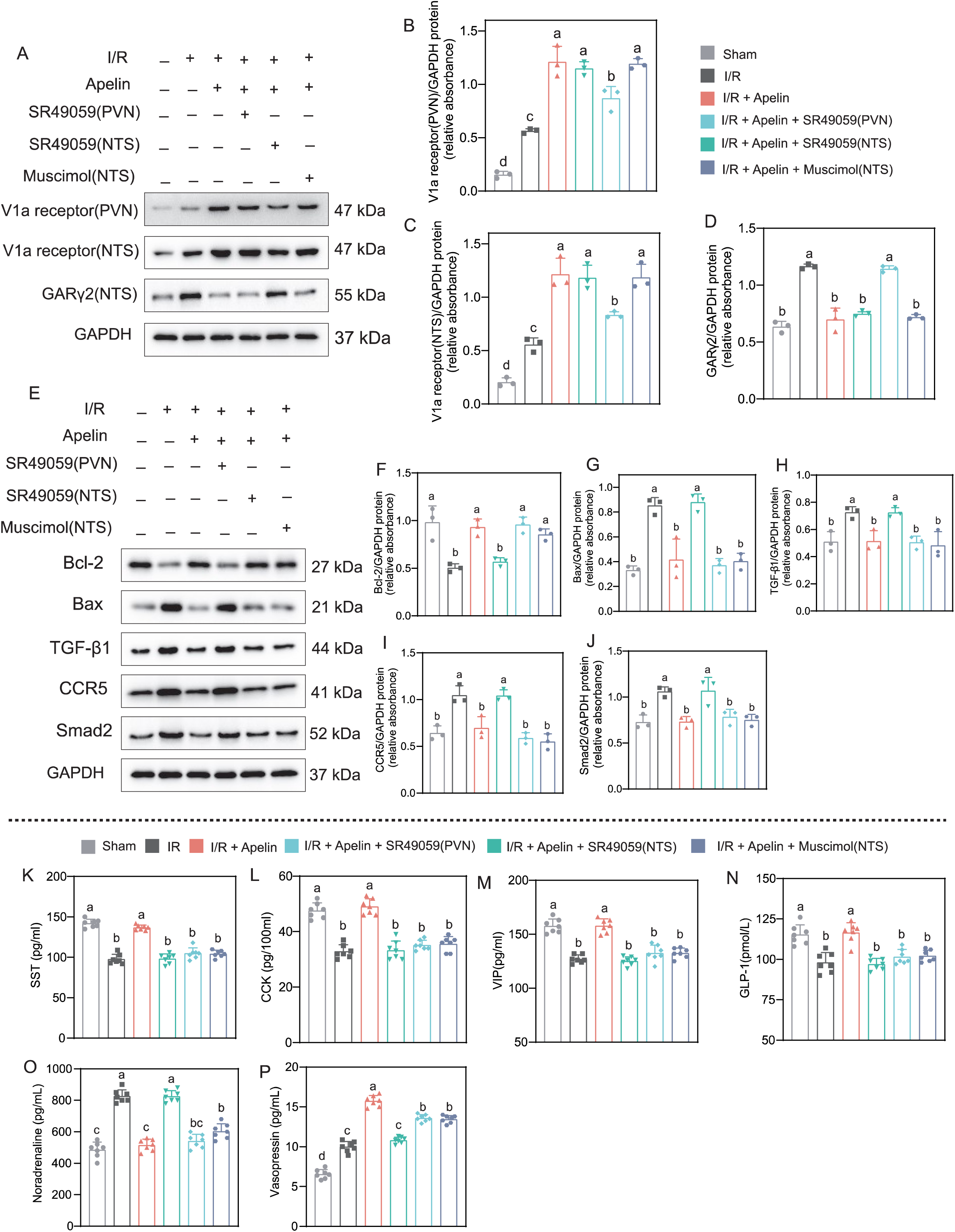
Mechanisms of effect of apelin-13 overexpression (AAV2-apelin-13 gene transfer) in the PVN on V1a receptor–mediated cardiac function improvement of I/R rats. **A** Representative WB for V1a receptor in the PVN or NTS and GARγ2 expression in the NTS, and **B–D** its quantification. **E** Representative WB for Bcl-2, Bax, TGF-β1, CCR-5, and Smad2 protein level in the heart and **F–J** its quantification. **K–P** The plasma levels of SST, CCK, VIP, GLP-1, noradrenaline, and vasopressin (n=7 in each group). Normality was tested using the Shapiro–Wilk test. Data (n=3 experiments) in **B**–**D** and **F**–**P** are shown as mean ± SD, and were analyzed using one-way ANOVA, corrected using Tukey’s multiple-comparisons test. In **B**–**D** and **F**–**P**, the differences are indicated by the abc annotation method.

## Discussion

This study provides novel evidence of neural regulation of cardiac functions via apelin-13 in the PVN upregulating V1a receptor in the PVN and NTS, and downregulating GARγ2 in the NTS. Effects of the V1a receptor antagonist and GARγ2 agonist furtherly prove that apelin-13 in the PVN is a novel pathway in centrally mediated neural control of myocardial injury, including MI and I/R rat models. Furthermore, myocardial ischemia–related apoptotic and inflammatory pathways are also involved in neural regulation of apelin-13 in the PVN.

First, in MI rats, apelin-13 expression was decreased in the PVN, but not in the NTS or RVLM. The PVN is a critical central regulator of autonomic and humoral responses. It receives afferent inputs from visceral receptors and circulating hormones, and then sends out projections to key cardiovascular brainstem centers and spinal cord preganglionic neurons, thereby modulating parasympathetic and sympathetic activity.^25^ The PVN is also involved in treatment of MI-induced heart failure by inhibition of oxidant signaling.^9^ Thus, we speculated that there were more pathways involved in cardiac function mediation by apelin-13 in the PVN. Interestingly, V1a receptor expression in the PVN and NTS was increased by apelin-13 microinjection into the PVN of MI rats, whereas GARγ2 expression in the NTS was reduced after apelin-13 microinjection, and these findings were furtherly proved in the I/R and MI models with apelin-13 gene transmission. In the RVLM, the pressor effect evoked by bilateral microinjection of apelin-13 as a modulating neurotransmitter in normotensive rats via V1a receptor affects vascular tone independent of GAR, while the presence of presynaptic V1a receptors affects vasopressin release from the PVN–RVLM projecting fibers.^17^ The observations in our study suggested that synaptic V1a receptor performed as communicator between the PVN and NTS for apelin-13–mediated nerve connections and GARγ2 expression in the NTS.

This study indicates that myocardium injury induces apelin-13 downregulation in the PVN and that upregulating apelin-13 in the PVN improves cardiac function. Thus, we speculated that the apelin/APJ system is involved in the PVN–NTS axis for cardiac function regulation, which is novel evidence of heart–brain interaction. The heart and the brain are linked by multiple feedback signals, and the concept of the cardio– cerebral syndrome in heart failure has been built on bidirectional interactions of failing heart and neuronal signals.^26^ A previous review article indicated that the mechanisms of brain–heart interaction include physiological effects of sympathetic and parasympathetic nerve activities, central pathways regulating the autonomic outflow, reflex control of the autonomic outflow, and integrative regulation of the autonomic outflow to the heart.^27^ More specifically, sympathetic and parasympathetic branches of the autonomic nervous system directly control the heart, and parasympathetic postganglionic fibers innervate the atrial and ventricular myocardium with releasing acetylcholine and VIP.^28^ Consistently, in our study, as critical mediator of synaptic inhibition in the brain,^29^ expression of γ2 subunit of GAR in the NTS was reduced by apelin-13 overexpression in the PVN, which indirectly elevated parasympathetic efferent excitability to increase PES.

V1a receptor is transmembrane protein that belongs to the superfamily of G-protein– coupled receptors. It has been demonstrated that the combined activation of V1a receptors by vasopressin in the PVN could elevate renal sympathetic nerve activity.^30^ Our data showed that V1a receptor expression in the PVN and NTS was increased in MI rats, and furtherly elevated by apelin-13 overexpression in the PVN of MI or I/R rats, demonstrating that V1a receptor is involved in apelin-13–mediated cardiac function.

The apelin/APJ system plays several important roles in neurohormonal regulation of heart function, including vasodilatory effect, positive inotropic effect, regulation of fluid balance, anti-apoptotic effect, and interaction with other neurohormonal systems.^31, 32^ From our results, the improved cardiac function mediated by apelin/APJ in the PVN involves PES increased by apelin-13 overexpression, including four effectors, namely SST, CCK, GLP-1, and VIP. SST has been demonstrated to exert a direct cardiocytoprotective effect against simulated I/R injury via SST receptor 1 and SST receptor 2 in cardiomyocytes and vascular endothelial cells.^33^ CCK, GLP-1, and VIP are involved in cardiovascular regulation with different mechanism in circulation.^34–36^ Thus, we speculate that PES plays an important role to achieve the heart–brain circuit.

Despite the meaningful findings, there were certain limitations to our study. Although we found novel evidence of neural control for cardiomyocyte injury, the mechanism interaction between the PVN and NTS is not fully clear, except for direct exposure to the neurotransmitter apelin-13 and vasopressin. Importantly, V1a receptor and GARγ2 in the NTS are indirectly regulated by the apelin/APJ system in the NTS, but postsynaptic GARs currents should be considered for modulating receptor expression. Therefore, there are some other unclear mechanisms involved in receptors interaction between the PVN and NTS. Upregulation of GARγ2 in the NTS or downregulation of V1a receptor in the PVN could not completely inhibit the function of apelin-13 in the PVN of MI rats, which indicates that the apelin/APJ pathway may play a leading role in neural regulation of cardiac function, but some other unclear mechanisms may also participate. Vasopressin present in the cerebrospinal fluid (CSF) possesses the ability to interact with the HPA axis. This interaction can enhance the release of adrenocorticotropic hormone from the anterior pituitary gland to modulate cardiac function.^37^ However, the levels of vasopressin in CSF were unfortunately not assessed. Finally, four peptides acting as neurotransmitters are released through a complex network of gut sensing and vagal mechanisms. Other neuropeptide levels could also be affected by apelin-13 in the PVN, however, these were not detected and explored in the present study.

## Conclusion

Apelin-13 in the PVN performs neural control for cardiac function via V1a receptor of the PVN and NTS, and GARγ2 in the NTS, as well as PES, offering novel evidence of brain–heart interactions.

## Acknowledgements

The authors thank LetPub (www.letpub.com) for linguistic assistance and pre-submission expert review; Bestcell (Hubei, China) for assistance with heart histology.

## Author’s contributions

W.Y., D.W., and XM.Z. conducted the experiments. CL.X. and W.Y. designed the experiments and analyzed the data. D.W. and CL.X. drafted the manuscript, and all authors edited the manuscript.

## Novelty and significance

### What Is Known?

- Currently, brain-heart crosstalk is known as neurocardiology, which focuses on the effects of brain injury on the heart as well as the impacts of cardiac injury on the brain.
- The apelin/APJ and GABAergic system are crucial pathways in the central nervous system that are involved in the neural control of the cardiovascular system within the hypothalamus.

### What New Information Does This Article Contribute?

- In the PVN, Apelin-13 upregulates the V1a receptor in both the PVN and NTS, while downregulating GAR γ2 in the NTS.
- The effects of the V1a receptor antagonist and GARγ2 agonist further demonstrate that apelin-13 in the PVN constitutes a novel pathway in centrally mediated neural control of myocardial injury.
- Four neuropeptides, namely somatostatin, cholecystokinin, glucagon-like peptide-1, and vasoactive intestinal peptide in serum are increased when apelin-13 is overexpressed in PVN to improve cardiac function.
- Myocardial ischemia–related apoptotic and inflammation pathway markers, including TGF-β1, CCR-5, Smad2, Bax, and Bcl-2, are involved in apelin-13 neurally mediated cardiac regulation.

